# *Identifying Refugia and Barriers to the Spread of* A. graminifolia *and* D. crumenatum *in Puerto Rico*

**DOI:** 10.1101/828517

**Authors:** Evan Foster, James D. Ackerman, Wilfredo Falcón L.

**Affiliations:** Department of Environmental Sciences, Colorado College; Department of Biology, University of Puerto Rico; Research Department, Eco-Caribe LLC

## Abstract

Establishment of new populations is contingent on overcoming abiotic and biotic barriers. While this applies to all species, these hurdles are at the forefront of invasion biology where prediction, prevention, eradication, and control strategies depend on an understanding and exploitation of barriers to establishment and spread. *Arundina graminifolia* and *Dendrobium crumenatum* are two invasive orchids spreading throughout Puerto Rico. Current records on their distributions across the island are sparse, and their interactions with the surrounding ecosystem are unknown. Through a direct population survey of all known localities, we identified a new, acquired enemy of both orchids: the orchid-specialist weevil, *Stethobaris polita*. In this study, we used niche modelling to identify suitable habitats for each orchid on the island and map their current distributions and interactions with *S. polita*, along with their distributions in the most extreme climate scenario in 2050, in order to contextualize projected patterns of establishment on the island. Our findings show that *D. crumenatum* flourishes in urban environments which also provide refugia from *S. polita*. In contrast, there is currently no refugia for *A. graminifolia* from *S. polita* attack, as it is more sensitive to the same climatic variables as *S. polita*. Furthermore, projections into the most extreme climate scenario suggests Puerto Rico will be unsuitable for *A. graminifolia* and *S. polita*’s survival, and become less suitable for *D. crumenatum*, by 2050.

## Introduction

Over the past century, our natural world has dealt with new, largely human-induced problems at an unprecedented scale. Terrestrial ecosystems are under siege from rapid deforestation and expanding urbanization, to a climate system shifting at a rapid rate, all of which are associated with greater frequencies of biological invasions (ref.). These changes are prevalent worldwide, and exacerbated on islands (D’antonio and Dudley, 1995; Denslow et al. 2009; Pulwarty et al. 2010) where some impacts are obvious and dramatic, whereas others are subtle, small and unexpected (e.g., Vitousek and Walker, 1989; O’Dowd et al., 2003; Sin et al., 2008; Recart et al., 2013). Certainly habitat heterogeneity and biotic and abiotic clines affect invasional success and geographic spread (Lockwood et al. 2007). For example, climate change can facilitate or impede invasive establishment and spread (Kolanowska et al. 2017; Ongaro et al, 2018). The heat island effect can increase surrounding temperatures, locally exaggerating current warming trends (Oke, 1973; Parker, 2009; McCarthy et al., 2010). Additionally, urban areas facilitate non-native prominence, and can provide refuge for invasive species from biotic resistance (McKinney, 2006; Burton et al., 2005; Holway, 1995; Vila & Pujadas, 2001), as is observed for the invasive orchid *Spathoglottis plicata* in Puerto Rico (Soifer and Ackerman, 2019).

In Puerto Rico, barrages of new, invasive species are being introduced (e.g., Rojas-Sandoval & Acevedo-Rodríguez, 2016; Falcón & Tremblay, 2018). Often overlooked are the increasing numbers of exotic orchids (Ackerman, 2007). Although orchids are disproportionately underrepresented among invasive species (Daehler 1998), those that are can have both positive and negative impacts (e.g. apparent competition, Recart et al. 2013). For this reason, it is important to identify recent invasions and factors associated with their establishment, spread and consequences to invaded communities.

In this paper, we focus on populations of *Arundina graminifolia* and *Dendrobium crumenatum*, two invasive Asiatic orchids in Puerto Rico. Current knowledge on both species in Puerto Rico, beyond herbarium collections, is rather sparse. The first naturalized population of *A. graminifolia* was discovered a few kilometers from an orchid nursery which had a stand of them on the nursery grounds, so it is likely they arrived via the horticultural trade as has happened elsewhere (Ackerman, 1995, 2012). Since their original discovery, both populations have spread across the island, as evidenced by local herbarium records. As both invasions continue to spread across the island over the coming decades, understanding their fullest possible extent ahead of time can provide useful information for future research and/or mitigation efforts on either species.

While modelling distributions and recording climatic responses, it is important to contextualize both invasions with their known ecological interactions and with the impending changes in the current climate system. For both orchids, there are no published records of acquired enemies (herbivores, seed predators) in Puerto Rico, but we have seen native *Stethobaris polita*, an orchid specialist weevil, attacking buds, flowers and fruits of both species. Such attacks on other orchids have led to severe flower damage, and their ovipositing in the fruits can compromise fruit set (Recart et al, 2013). The implications of this interaction for both species on the island remains unknown, however the introduction of additional exotic orchids might pose additional risk to native orchids through apparent competition (Recart et al, 2013). In this way, understanding the extent of these interactions with *S. polita* across the island can also be particularly useful in understanding their greater ecological impacts. Previous research suggests *S. polita* does not inhabit urban landscapes, either due to an inability to penetrate or to establish within such environments (Soifer & Ackerman, 2019). For orchids like *D. crumenatum* that have naturalized within these areas, they can be freed from such herbivore attack. Such refugia has been observed in *S. plicata*, another exotic orchid in Puerto Rico (Soifer & Ackerman, 2019). *A. graminifolia* is found at higher elevations than the current urban extent, however there might be additional climatic barriers providing refugia from *S. polita* attack.

Both species exhibit signs of high climate sensitivity, making their responses to climate change particularly important in modelling their distributions. *A. graminifolia* is found naturally at higher elevations, indicating a sensitivity towards higher temperatures. Previous research has established a negative relationship in *A. graminifolia*’s habitat suitability globally under any climate change scenario, although the specific implications for this loss is unknown for Puerto Rico (Kolanowski and Konowalik, 2014).

For *D. crumenatum*, there is no standing research on its limiting climatic variables, much less it’s response to climate change. However, a strong --and perhaps unusual-- climatic response might be expected for *D. crumenatum*, since it’s reproductive success and population health directly relies on the weather. The gregarious flowering of *D. crumenatum* triggered by rain-induced temperature drops have been well-documented (Seifriz, 1923; Goh et al. 1982; Brooks and Hewitt, 1909). In order for current distribution models of *D. crumenatum* to remain applicable in the coming decades, they must account for scenarios with an altered climate.

In general, little to no research has been conducted on the distributions of *A. graminifolia* and *D. crumenatum* in Puerto Rico, or how these populations relate to their greater environment. As both populations continue to spread and establish on the island, it is important to understand how enemy acquisition, urbanization, and climate change will influence both invasive populations in the future.

### Objectives

Our research intends to establish an understanding of the variables influencing *A. graminifolia* and *D. crumenatum* distributions in Puerto Rico in the present, allowing us to predict what their futures hold on the island. In order to do so, our project is split into three objectives, each probing a variable that might have a demonstrable effect on both/either orchid species. Our first objective is to determine whether enemy release or enemy acquisition is occurring for *A. graminifolia* and *D. crumenatum*, and to what extent it may affect reproductive success. Our second objective is to assess whether anthropogenic disturbances provide refugia for both orchid species from herbivore attack. Our third objective is to determine the responses of *A. graminifolia* and *D. crumenatum* to future climate conditions; primarily along the basis of their distributions and relationships.

### Hypotheses

#### Enemy Acquisition

The orchid-specialist weevil, *Stethobaris polita*, has been observed feeding on the reproductive structures of both orchid species (field observation). However, there are other herbivorous invertebrates that attack orchids in Puerto Rico, such as other weevils, aphids, and snails (summarized by Martorell and Gaud, 1974). It can be expected that some of these other herbivores will also attack *A. graminifolia* and *D. crumenatum*, although the prevalence of these interactions is still unknown. Regardless, if *S. polita* frequently attacks *A. graminifolia* and *D. crumenatum*, then we expect that other herbivores will also attack them. In quantifiable terms, we expect areas with higher abundances of enemies to subsequently have reduced reproductive success, indicated by lower fruit production and/or pollinarium removals.

#### Presence of Refugia from Enemies

Populations of *D. crumenatum* can be found in the urban landscapes of San Juan. Most likely, these orchids were originally planted artificially by humans, where they then had reproductive success and naturally established within urban forests and other urban areas (personal observation). At the same time, a common culprit of orchid herbivory, the orchid-specialist weevil *S. polita*, is unable to travel into or establish within urban areas, remaining largely absent from those locations (Soifer and Ackerman, 2019). If there is this disparity between herbivore and plant, then there will be a forest-urban gradient for the *D. crumenatum*, similar to what is seen in the S*. plicata*. At the same time, however, since *A. graminifolia* has naturalized only at elevations above 500 m, far above the extent of most current urban areas, the reliance of *Arundina* on the bioclimatic variables associated with higher altitudes will prevent the species from establishing within lowland urban centers. Furthermore, the presence of *A. graminifolia* within those areas are completely facilitated by/ reliant upon human dispersion and care. However, other barriers to enemy establishment that might be associated with the current extent of *A. graminifolia* (e.g. bioclimates, soil types, elevation) could create refugia for *A. graminifolia* from its enemies.

#### Response to Climate Change

Previous models of *A. graminifolia* in Central and South America show a noticeable decrease in suitable habitats under every climatic warming scenario provided by the IPCC (Kolonawska and Konowalik, 2014). This makes sense, as *A. graminifolia* are only found at higher elevations in Puerto Rico, indicating a possible reliance/preference on the cooler bioclimatic variables associated with those elevations. This fragility to warmer temperatures might force an altitudinal range shift to higher elevations (Lenoir et al. 2008; Walther et al. 2002). Clearly, if invasive populations of *A. graminifolia* are vulnerable to future climate change, then we would expect either a future die-off, or a restricted growth in distribution, of the invasive *A. graminifolia* populations in Puerto Rico within the coming decades, possibly minimizing their overall effects on the native species in Puerto Rico.

On the other hand, *D. crumenatum* relies almost exclusively on a surrounding temperature drop to signal a synchronous flowering (Seifriz, 1923; Goh et al. 1982; Brooks and Hewitt, 1909). Considering how the orchids rely on temperature change, rather than a specific temperature, the vulnerability of such orchids to climate change will largely be determined by the temperature variability at lower altitudes as a result of climate change. *Dendrobium crumenatum* could hypothetically see greater reproductive success and more frequent flowering, or the exact opposite. Regardless, the distribution of *D. crumenatum* in Puerto Rico will be affected by climate change in a major way.

## Materials and Methods

### Study Site

The study was conducted during June and July 2019 across the Carribean island of Puerto Rico. Populations were monitored mainly along roadsides and in front yards, although a few populations were found along trails and streams. The locations of individual populations were identified from herbarium collections and personal communications. Populations were also identified through incidental discovery, either during travels between known populations, or through intentional scouting in previously unexplored municipalities.

We collected data from the following municipalities: Adjuntas, Aibonito, Barranquitas, Caguas, Carolina, Cayey, Cidra, Jayuya, Lares, Luquillo, Naranjito, Orocovis, Patillas, Río Grande, San Juan, San Sebastian, Trujillo Alto, and Utuado.

### Field Work

We collected GPS coordinates and plant data for populations along roadsides, private properties, and trails. Populations were only considered if there was evidence of naturalization and/or recruitment. Populations could meet this criteria in a number of ways: the presence of reproductive structures (i.e. fruiting bodies) on at least one plant, the presence of recruits in the population, or sufficient circumstantial evidence that the population was not cultivated. Even if a population was originally cultivated, it was still included if successful recruitment was evident, acknowledging the role of the horticultural trade as a vector in exotic orchid dispersal.

At each designated study site, we identified a “target plant,” which we categorized as an orchid with reproductive structures closest to the center of the population. We collected the waypoints for the target plant with a GPS system, and measured the distance of the target plant to its five nearest neighbors (within a 30 m radius). We averaged these distances to calculate a proxy for population density. For plants without reproducing neighbors within a 30 m radius, their density measure is greater than or equal to 30 m (i.e. the average of nearest neighbors is at least 30 m).

At each target plant and their five nearest neighbors, we gathered data on its anatomical and biological characteristics, particularly those that are important for establishment and success in the next generation. More specifically, for populations of *D. crumenatum* we measured the number of active shoots, the length and width of the largest pseudobulb, plant height from ground, total number of fruits, and the number of damaged fruits. Due to the erratic flowering of *D. crumenatum*, flower data could not be taken at most sites. When plants were found in flower, we counted total flower number and identified reproductive success by identifying pollinaria removals and pollinations. Due to the large number of flowers associated with the synchronous flowering spectacle of *D. crumenatum*, we sampled all available flowers or a random sample to a maximum of 30 flowers for pollinarium removals and pollinations.

For *A. graminifolia*, we measured the petal width, throat width, lip length, and height of each plant's flowers. We measured the height of fruiting bodies. We also recorded the number of scars for each active shoot. Similar to *D. crumenatum*, we identified pollinaria removal and deposits. We recorded the damage to flower petals, throat, lip, and sepils, along with the damage to fruits.

At every population, we recorded the number of *S. polita* present, along with the presence of other possible enemies. We categorized weevil presence at a waypoint from either weevil sightings or evidence of weevil damage and damaged reproductive structures (buds, flowers, fruits). We quantified the degree of damage to plants and populations by designating a damage score for each flower and fruit, on a 1-10 scale. By scoring each structure individually, we could effectively average the total damage to each plant and to entire populations.

### Distribution Modelling and Mapping

Land use will be quantified into three categories of disturbance, similar to previous studies: urban forests (0-33% undisturbed habitat), rural forests (34-66% undisturbed habitat), and continuous forest (67-100% undisturbed habitat) (Soifer and Ackerman, 2019). Specific land cover type and habitat composition of our study sites was determined using QGIS and the land cover map from the Puerto Rico GAP Analysis Project (PRGAP) (2006).

### Maxent

We modelled the distributions of *A. graminifolia* and *D. crumenatum* using Maxent v 3.4.1 (Phillips et al. 2019), similar to previous studies on invasive species distribution in both the present and future (Soifer and Ackerman, 2019; Kolanowska and Konowalik, 2014). We ran the presence-only data for both species against ten environmental layers, including eight climatic layers from Worldclim Version 2––Global Climate Data (Fick and Hijmans, 2017). The climatic layers chosen were based off of similar studies on the same study species, or within the same study area, which we hope to compare our data the closest to: annual mean temperature (Bio 01), temperature seasonality (Bio 04), maximum temperature of warmest month (Bio 05), minimum temperature of warmest month (Bio 06), mean temperature of driest quarter (Bio 09), precipitation seasonality (Bio 15), precipitation of wettest quarter (Bio 16), and precipitation of driest quarter (Bio 17) (Soifer and Ackerman, 2019; Kolanowska, 2014; Recart et al., 2013). This set of environmental layers will be narrowed down before publication, as we have not run GLM’s between each climatic layer to eliminate those that are highly correlated. We also included land cover and elevation (PRGAP 2006). Bioclim variables and elevation were set at a spatial resolution of 30 arc-seconds (~1 km), with land cover at 15m² (Gould et al., 2008). We prepared the layers for Maxent on ArcGIS to match the extent, cell size, and coordinate system of the PRGAP land cover map. We also used ArcGIS to match our GPS coordinates with the coordinate system of the environmental layers.

Samples included a total of 30 populations of *D. crumenatum*, 28 populations of *A. graminifolia*, and 140 populations of *S. polita*. For any cell with more than one population, only one was used to contribute to the model. Since presence points for *S. polita* would be limited to the presence points of our two orchids, their modelled distribution might create an incomplete picture of their overall distribution. To counteract this discrepancy, we combined our *S. polita* presence points with those recorded with a third orchid, *Spathoglottis plicata* (see: Soifer and Ackerman, 2019). For this reason, the number of populations of *S. polita* is noticeably higher than those of either orchid.

We ran logistic models for *A. graminifolia*, *D. crumenatum*, and *S. polita* under the default settings, while running a 10-fold cross-validation to estimate error around the mean model, which has been identified as a reliable method for projecting onto novel climate scenarios (Elith et al 2010). We applied the threshold rule of Equal Training Sensitivity and Specificity, similar to previous studies.

We used these same climatic layers as we project orchid population response under various climatic scenarios (RCPP 2.6 and 8.5), supplied by Ramirez and Jarvis (2008), and utilized by similar studies (Coops and Waring, 2011; Kolanowska and Konowalik, 2014; Barredo et al. 2015; Kolanowska et al. 2017). While the accuracy of such projections remain unknown, and their overall reliability remains highly contested (Araújo and Peterson, 2012), we are following the guidelines set by previous studies to ensure more accurate modelling (Elith et al 2010).

### Statistical Analysis

Along with modelling species distributions, Maxent also provides multiple statistical tests on model significance and performance. We used the area under the receiver operating characteristic (ROC) curve (AUC) to assess model quality and discern model output from a random prediction. We conducted jackknife tests for each important variable, allowing us to determine the relative importance that variable has in determining the model.

Following the conclusion of this REU, we will run chi squared tests and generalized linear models for abundance of herbivores on the degree of herbivory, reproductive success (fruit set and production), and plant density of *A. graminifolia* and *D. crumenatum*. We will also run abundance of herbivores against the flowering of *Arundina*, along with the relationship between *Arundina’s* flower size and fruit set. We These will all be conducted with climatic variables held constant. Together, these tests should illustrate which factors are most important in explaining various outcomes of orchid distribution and health.

## Results

### Enemy Acquisition

Field observations show that both *A. graminifolia* and *D. crumenatum* have acquired a new enemy in the form of *S. polita*. Enemy attack is more common for *A. graminifolia* than it is for *D. crumenatum*. Of the 30 populations of *D. crumenatum* surveyed, only 6 contained evidence of *S. polita* damage. In comparison, 20 of the 28 *A. graminifolia* populations surveyed had evidence of *S. polita* damage. Some *A. graminifolia* flowers could be seen with damage to their sepils, lip, and/or petals, with *S. polita* present (Fig. 6). *S. polita* was also seen boring into *A. graminifolia* fruits. Identical damage markings were found on other *A. graminifolia* flowers and fruits, even if no *S. polita* were present, indicating previous attack. Observations of *S. polita* on *D. crumenatum* were limited to *D. crumenatum*’s flowering events, with similar damage to *D. crumenatum*’s flowers as observed on *A. graminifolia* (Fig. 6). Ovipositing holes in *D. crumenatum’*s fruits were observed as well.

### Distribution Modelling and Mapping

The average AUC values for each species indicate good model fit (*A. graminifolia*: AUC = 0.934, standard deviation = 0.049; *D. crumenatum*: AUC = 0.911, standard deviation = 0.051; *S. polita*: AUC = 0.884, standard deviation = 0.057) (Fig. 2). After applying an equal training sensitivity and specificity threshold, there were variations in the average presence threshold for *A. graminifolia*, *D. crumenatum*, and *S. polita* (0.2131, 0.3216 and 0.1797 respectively).

For *A. graminifolia*, Maxent predicts suitable habitat within the higher elevations of the Cordillera Central, Carite Forest and El Yunque National Forest, which matches our field observations of population occurrence (Fig. 1C).

For *D. crumenatum*, our models show an affinity towards areas of high land use, with suitable habitat coinciding with urban areas, particularly San Juan and the Caguas-Cayey region, which is where a high density of *D. crumenatum* populations were observed (Fig. 1B). *Dendrobium crumematum* appears to prefer lower elevations and largely disturbed habitat, with it’s extent noticeably excluding El Yunque National Forest and Cordillera Central. Areas of high land use are not the sole variable necessary for survival. The dry forests on the South side of the island remain unpreferable, even with large urban areas like Ponce.

For *S. polita*, our models show a negative relationship with land cover. The intact forests of El Yunque and Carite are deemed the most suitable habitat, with hot spots projected in those areas (Fig. 1A). At the same time, urban areas are almost entirely excluded from *S. polita*’s extent, particularly San Juan and Caguas, which matches field observations. Additionally, *S. polita* appears to prefer higher elevations in wet conditions, with their range extending along the Cordillera Central, except for portions with the highest elevations, and into the northern karst region. The scattered pattern across the Northern Karst region are likely a product of the microclimate conditions from the uneven elevation changes of the magotes.

**Figure 1A:**
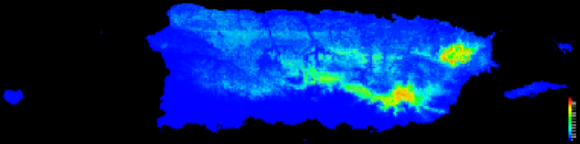
Average Maxent output of habitat suitability for *S. polita* under a current climate scenario. Warmer colors indicate areas with better predicted conditions for survival. Most notably, the model suggests preferable conditions in the intact forests of El Yunque National Forest and Carite Forest Preserve, while developed areas are considered unpreferable.

**Figure 1B:**
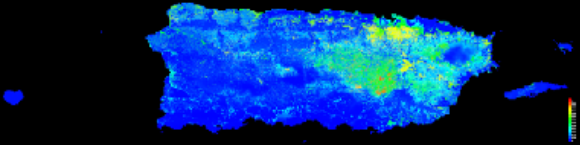
Average Maxent output of habitat suitability for *D. crumenatum* under a current climate scenario. Warmer colors indicate areas with better predicted conditions for survival. The model suggests *D. crumenatum* can thrive in areas of high land use, particularly in San Juan and the Caguas-Cayey region. However, urban areas in Puerto Rico’s dry forests on the South side of the island remain unpreferable, due to other climatic factors.

**Figure 1C:**
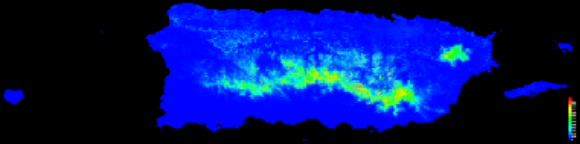
Average Maxent output of habitat suitability for *A. graminifolia* under a current climate scenario. Warmer colors indicate areas with better predicted conditions for survival. The model suggests that the high elevations of mountainous regions in the center of the island provide the best conditions for *A. graminifolia*.

**Figure 2:**
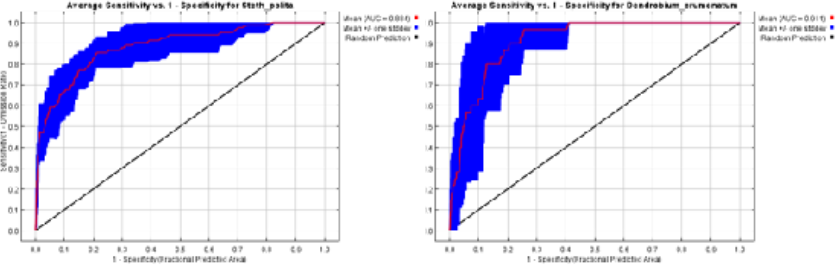
The Area Under the Receiver Operating Characteristic (ROC) curve (AUC) for each species over 10 replicated runs. An AUC > 0.5 represents a non-random prediction, while AUC > 0.80 represents a good model and AUC > 0.90 indicate a very good model. However, these thresholds are arbitrary above random and should be assessed on a per-model basis. A) ROC curve for *A. graminifolia*; AUC = 0.934, standard deviation = 0.049. B) ROC curve for *D. crumenatum*; AUC = 0.911, standard deviation = 0.051. C) ROC curve for *S. polita*; AUC = 0.884, standard deviation = 0.057.

Land cover is one of the most important variables in determining the distribution of all three species (Fig. 3). In each jackknife test, training gain is lowest when land cover is removed from the model, indicating that it contains information found in no other variables (Fig. 3). Land cover also gives *D. crumenatum* the highest training gain in isolation, providing the most useful information on its own. Interestingly, training gain is higher for *D. crumenatum* with land cover in isolation than when only land cover is removed from the model. Other than land cover, precipitation and temperature seasonality, along with precipitation during the driest quarter contribute to the model (Fig. 3). This makes sense, considering *D. crumenatum*’s reliance on rainfall-induced temperature drops to trigger their gregarious flowering. For *A. graminifolia* and *S. polita*, Max. Temperature in the Warmest Month gives the highest training gain in isolation, with all other temperature layers and elevation providing similar training gains in isolation (Fig. 3). Elevation and temperature are likely closely related, with altitudinal preference possibly a product of their sensitivity to temperature conditions. Correlation tests still have to be run to identify highly correlated variables. Clearly, both species are sensitive to temperature conditions.

**Figure 3:**
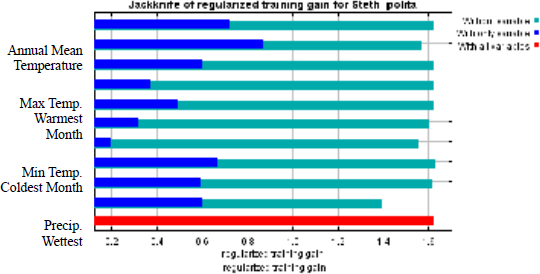
Jacknife tests calculate the regularized training gain when each variable is removed from the model and when only one variable is included in the model. Comparing training gains between variables can identify variable importance. For all three species, training gain is lowest when Land Use is removed from the model, indicating that it contains information found in no other variables. For *A. graminifolia* and *S. polita*, Max. Temperature Warmest Month gives the highest training gain in isolation, while Land Use gives *D. crumenatum* the highest training gain in isolation. These variables provide the most useful information on their own.

Co-occurrence maps reveal how variations and similarities in response to the selected environmental variables influence the degree of inter-species interactions. *D. crumenatum* heavily favors towards areas of high anthropogenic disturbance, while *S. polita* heavily excludes such environments. This division leads to sparse overlap in their extent, and limited areas of co-occurrence (Fig. 5). In comparison, the similar response of *A.graminifolia* and *S. polita* to environmental variables results in nearly all of *A. graminifolia*’s extent to coincide within *S. polita*’s (Fig. 5).

Response to climate change follows the same species-level patterns observed in variable importance and habitat suitability. Under the most extreme climate change scenario (RCP8.5), our model suggests Puerto Rico will become uninhabitable for both *A. graminifolia* and *S. polita*, with no portions of the island projected as any degree of suitable (Fig. 4). At the same time, our model predicts that such climate conditions will create slightly less suitable conditions for *D. crumenatum* in Puerto Rico (Fig. 4). Current suitable habitats will become less suitable, while the total area of suitable habitat for *D. crumenatum* on the island will noticeably decrease (Fig. 4). However, *D. crumenatum* is still expected to remain on the island, particularly within urban centers. This trend is not particularly strong, since the maximum distribution modelled for *D. crumenatum* actually shows an increase in habitat suitability and geographic extent. However, the overall trend of decreased suitability remains.

**Figure 4:**
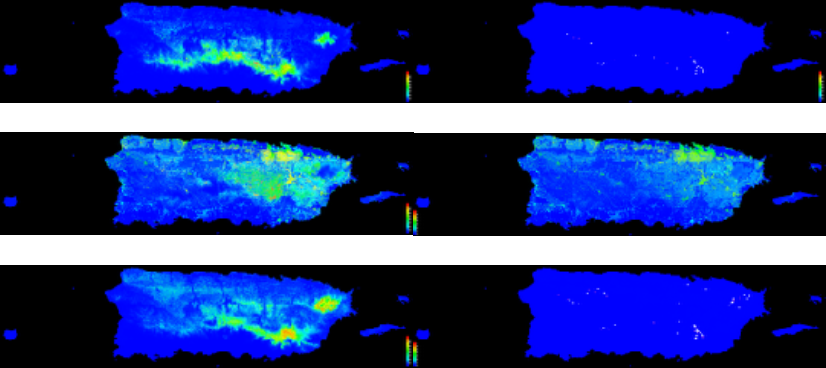
Compares the average Maxent output of habitat suitability for each species in the current climate (left) against the habitat suitability in 2050 with the most extreme climate scenario (RCP8.5) (right). Warmer colors indicate areas with better predicted conditions for survival.

**Figure 5:**
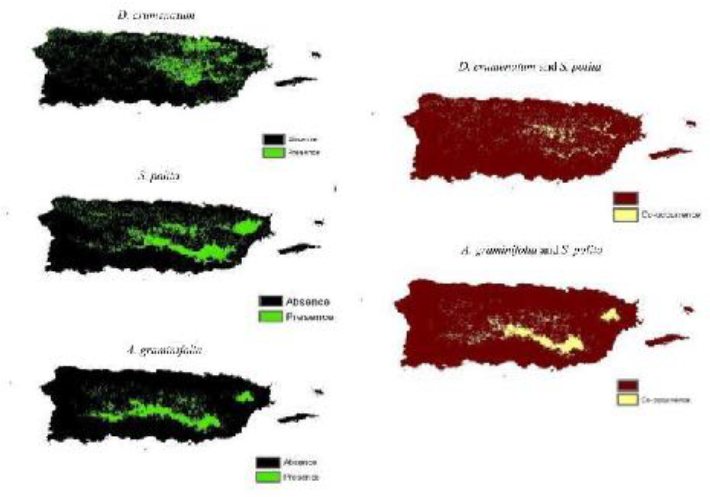
Maps on the left show the distribution of all three species after an Equal training specificity and sensitivity threshold was applied. Green represents presence, while black shows absence. Threshold values for A. graminifolia = 0.21; D. crumenatum = 0.32; S. polita = 0.18. Maps on the right combine the presence-absence maps of each orchid to S. polita’s. Yellow areas indicate regions where both species co-occur.

**Figure 6:**
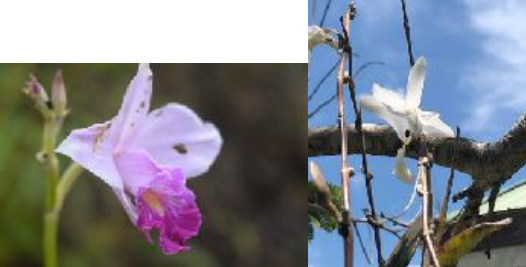
Photos taken by authors of *A. graminifolia* and *D. crumenatum* with petal damage from S. polita attack and a resident *S. polita.*

## Discussion

### Factors Driving Orchid Distributions and Biotic Interactions

Accessibility and sensitivity to areas of high anthropogenic disturbance, along with comparisons between determining climatic factors, are what define the distributions of *A. graminifolia*, *D. crumenatum*, and their biotic interactions with *S. polita*.

Land cover is the factor that is most associated with the distribution of *D. crumenatum* across Puerto Rico. The overwhelming prevalence of naturalized *D. crumenatum* within urban areas is a confluence of multiple factors, which allowed for a rapidly spreading *D. crumenatum* population. First and foremost is the integral role the formal and informal horticultural trade has had on dispersing *D. crumenatum* to gardens and urban areas across the island. Although it is unknown how *D. crumenatum* was first introduced to Puerto Rico, a majority of the world’s naturalized plants are deliberately introduced, particularly through the horticultural trade (Mack and Erneberg, 2002; Mack, 2003), and the more popular a plant is, the greater the propagule pressure and the higher the probability of invasional success (Dehnen-Schutz et al., 2007). The rapid spread of *Dendrobium crumenatum* across the island over the past decade is certainly connected to its increasing popularity.

Since *D. crumenatum* is such a recent invasion/naturalization, the evidence of recruitment from initial cultivation is profound. Almost all naturalized populations of *D. crumenatum* appeared to be sourced from a nearby cultivated parent plant. Many orchids were planted in front yards along roadsides, and the role of roads as an effective corridor for exotic seed dispersal has already been identified (Parendes and Jones, 2000; Mortensen et al, 2009). Rivers are also likely corridors for *D. crumenatum* spread; however, thus far we are aware of only two populations along riparian habitats.

Furthermore, the current urban extent coincides well with the expected climatic preferences of *D. crumentatum*, as they prefer lower elevations in moist to wet environments, which matches the climatic conditions of urban areas such as San Juan and Caguas (Ewel & Whitmore 1973; López-Marrero & Villanueva Colón 2006). The reliance of *Dendrobium crumenatum* on rain-induced temperature drops to trigger mast flowering creates expected effects on its distribution. Areas with lower amounts of rain at less frequent intervals, such as the dry forests along the southern coast, are uninhabitable for *D. crumenatum*. Our analyses indicate that temperature and precipitation seasonality likely play a role in their establishment, and response curves indicate that the lower threshold of precipitation during the year also plays a role. Further extrapolation of the response curves still needs to be conducted to determine the specific responses *D. crumenatum* has to these variables.

Land cover provides widespread refugia for *D. crumenatum* from *S. polita* attack, adding another complementary aspect in facilitating *D. crumenatum* establishment within those areas. Exotic increase and native decrease towards urban centers is a widely observed phenomenon (Burton et al. 2005; Almasi, 2000; McKinney, 2002). Our findings reaffirm *S. polita* absence from the urban matrix (Soifer & Ackerman, 2019). With a majority of *D. crumenatum* residing almost entirely within areas of high land use, there is little co-occurrence with *S. polita* and limited biotic interaction (Fig. 5), although that does not exclude the possibility that, were it not for land cover, the biotic interaction could noticeably increase. Land cover alone cannot explain species naturalization, particularly for an invasive orchid Such tests will be carried out in the upcoming weeks.

In comparison, no environmental refugia was identified for *A. graminifolia* from *S. polita* attack, with almost all projected presence points falling within *S. polita*’s extent (Fig. 5). There are key differences between *A. graminifolia* and *D. crumenatum* that distinguish their ability to take full advantage of possible refugia and characterize their distributions.

Unlike *D. crumenatum*, *A. graminifolia* appears highly sensitive to all temperature variables. This sensitivity to high temperatures throughout the year results in establishment only within high elevations, with a minimum benchmark around 500 m. This lower altitudinal limit prevents *A. graminifolia* from spreading downwards towards the urban extents of San Juan and Caguas. For one *A. graminifolia* intensively cultivated at lower elevations, we observed significant heat damage on the flowers.

Lack of access to areas of high land use is not the only factor preventing *A. graminifolia* from escaping weevil attack. Our model suggests that the distribution of *A. graminifolia* is decided by the same environmental variables as *S. polita*, but demonstrates a greater sensitivity to each (Fig. 3). In effect, there should be little to no climate conditions where *A. graminifolia* could survive and *S. polita* could not.

The previously stated role of roads as a corridor for exotic seed dispersal is evident in the spread of *A. graminifolia* across the island. Due to the longer residency on the island, *A. graminifolia* has infiltrated protected forest reserves far removed from human cultivation, with populations thriving alongside the steep, open-canopied slopes bordering their roads.

Although human dispersal and the proximity to roads likely play a similar role to facilitating the spread of *A. graminifolia* across the island, highly disturbed areas of land cover might also hinder *A. graminifolia* establishment within those areas. Since *A. graminifolia* is a terrestrial orchid, it is likely far more difficult for its seeds to germinate and for successful recruitment to take place within the urban concrete jungle than it is for the epiphytic *D. crumenatum*, which can easily establish on the multiple cultivated trees present in urban areas, pending the presence of the necessary mycorrhizal fungi.

Additionally, unlike *D. crumenatum*, the orchid resource base within Puerto Rico’s mountain towns might be too sparse to support *A. graminifolia*’s acquired pollinators, the *Centris haemorrhoidalis* and *Apis mellifera* bees, at a population level that would optimize their pollination chances. Pollination surveys of *A. graminifolia* show roughly 11.4% of *A. graminifolia* flowers might experience a pollinator visit in a rural environment bordering El Yunque (Ackerman et al., unpublished). Indeed, land cover might exacerbate the infrequency of pollination for *A. graminifolia*, which relies on deception rather than rewards, like *D. crumenatum*. This might explain why, despite frequent cultivation as an ornamental within developed areas, most *A. graminifolia* plants within those environments are without fruits or signs of recruitment, while *D. crumenatum* has ample examples of both. Another possibility is that the terrestrial nature of *A. graminifolia* increases its risk of human intervention after recruitment. While *A. graminifolia* are frequently used as ornamentals and are often planted in visually-appealing patterns (i.e. a straight line lining a fence or a house), their wind-dispersed seeds rarely abide by these pre-arranged designs. It is possible that *A. graminifolia* recruits are removed at a much higher rate than *D. crumenatum* recruits, whose gradual spread on surrounding trees is unintrusive to their growers’ daily routine or garden upkeep. Surveys with orchid growers would have to be conducted to formally prove that there is a difference in the rate of human intervention of ornamental recruitment between terrestrial and epiphytic plants.

Our projections of *A. graminifolia* largely coincide with the previous models of their distribution on the island, despite the addition of multiple new presence localities between both models (Ackerman, unpublished), further supporting the validity of Maxent to accurately predict species distribution with small sample sizes.

Additionally, our models of current distribution agree with those within global *A. graminifolia* models (Kolanowska & Konowalik, 2014). While Maxent cannot accurately predict distributions in one locality from another (Ackerman, unpublished), it can accurately predict habitat suitability in new areas by extracting data from multiple localities across the entire climatic niche.

### Distributions and Interaction Response to Climate Change

Models of future climate change in Puerto Rico predict an above average temperature gain and decreased trends in precipitation, coinciding with more intense droughts (Khalyani et al. 2015). Further analysis and extrapolation needs to be conducted on the model used from worldclim to determine whether it’s predicted conditions match the observed changes and GCMs. Until then, the preliminary assumption is that they remain consistent with those trends.

Within this context, Puerto Rico would be void of all suitable habitat for *A. graminifolia* and *S. polita*, and neither species are expected to be present (outside of cultivation) on the island by 2050 in the most extreme climate scenario, while *D. crumenatum* will see only a slight reduction in its extent on the island.

The dramatic response of *A. graminifolia* to drastic climate change is not surprising. It is globally projected that such a climate scenario would reduce the range of suitable habitats within the invasive range of *A. graminifolia* by 88% by 2080 (Kolanowska & Konowalik, 2014).

In comparison, the less dramatically reduced extent of *D. crumenatum* represents the diversity in orchid response to climate change that previous studies have observed, with some orchids facing extinction while others proliferate (Kolanowska et al., 2017; Ongaro et al, 2018). So far, orchid response to climate change remains specific to each species, as two orchids with similar climatic preferences can respond to climate change differently (Kolanowska et al., 2017).

With land cover held constant across future projections, the decreased but remaining suitability for *D. crumenatum* within those areas suggests a heavy facilitation for survival by land cover. While it would be expected for a species like *D. crumenatum*, which prefers the warmer temperatures of lower elevations, to shift or expand its range towards higher elevations in a warming climate, the projected decrease in precipitation across the island appears to prevent such a transition. Low precipitation already restricts *D. crumenatum* from suitable temperature conditions on the island, as the dry forests of Puerto Rico remain uninhabitable despite similar temperatures and elevation as San Juan. The remaining presence within urban centers should bring cause for concern, as it would further its prominence within human centers, making it a highly noticeable and common “weed.” In order to prevent the spread of *D. crumenatum* in Puerto Rico, measures must be taken soon to control the population and regulate its dispersal in the horticultural trade. The areas highlighted in this paper can provide a useful map for their management. Management of *A. graminifolia* might not be necessary, or not as urgent as it is for *D. crumenatum*. If anthropogenic-climate change will manage the population in Puerto Rico, any funding or efforts might prove unnecessary.

One of the most important factors in determining a species’ response to climate change is not necessarily a climatic variable. Often forgotten in SDM’s projecting future distributions are the possible changes in biotic interactions (Tylianakis et al. 2008; Gilman et al. 2010; Davis et al. 1998). For *A. graminifolia* and *D. crumenatum*, their future distributions cannot fully be understood without an analysis of their biotic interactions during those times. Although the extent to which *S. polita* attack effects either plant is still being analyzed, its projected absence within a future climate scenario might implicate the extent of *D. crumenatum*, which is projected to supposedly survive drastic climate change by 2050. However, the projected extent of *D. crumenatum* within that climate scenario makes weevil absence seemingly inconsequential, as their projected extent within urban environments already excludes weevil presence. It is possible, however, that the more dramatic response of *S. polita* to climate change might provide an RCP scenario where it experiences a reduction in suitable habitat, such as an altitudinal range shift, freeing *D. crumenatum* from any enemy attack and its possible effect of biotic resistance. However, such a scenario could also see *D. crumenatum*’s suitable habitat extending to higher elevations, thereby negating the aforementioned release from enemy attack. Such interactions will be extensively explored as our data is run against the remaining RCP pathways.

Such freedom from enemy attack due to less drastic climate change seems unlikely for *A. graminifolia*, since it appears as sensitive, if not more, to projected climate change as *S. polita*. In this way, no climate scenario should offer *A. graminifolia* any form of environmental refugia from *S. polita*. We plan to test this theory over the coming weeks, as we run both species against lower RCP pathways.

These differences between *D. crumenatum* and *A. graminifolia* in their changing interactions with *S. polita* under climate change can be extrapolated across all orchids in Puerto Rico subject to *S. polita* attack. Orchids more accustomed to lower elevations and resilient to increasing temperatures could escape *S. polita* attack, while those found at higher elevations and are less resilient to increasing temperatures and/or increased droughts will likely face a similar fate to that of *S. polita* in the wake of impending climate change.

In order to fully understand the distributions of both orchid species in the future, similar population surveys must be conducted on their other biotic interactions. Niche modelling of both orchids’ pollinators will shed significant light on how either orchid will truly respond to climate change in the near future, particularly under more conservative RCP pathways.

